# Leveraging a hybrid cross-disciplinary training model to accelerate global bioinformatics capacity

**DOI:** 10.64898/2026.01.21.700760

**Authors:** Taras K. Oleksyk, Daryna Yakymenko, Sylwia Bożek, Viorel Munteanu, Wojciech Pilch, Zoia Comarova, Victor Gordeev, Grigore Boldirev, Dumitru Ciorbă, Viorel Bostan, Christopher E. Mason, Alexander G. Lucaci, Nadiia Kasianchuk, Daria Nishchenko, Victoria Popic, Andrei Lobiuc, Mihai Covasa, Martin Hölzer, Joanna Polanska, Alex Zelikovsky, Vasili Braga, Mihai Dimian, Paweł Łabaj, Serghei Mangul

**Affiliations:** Department of Biological Sciences, Oakland University, Rochester MI, USA; NGO “Bioinformatics for Ukraine”; Department of Biology and Department of Biological Sciences, Uzhhorod National University, Uzhhorod, 88000, Ukraine; Department of Cell Biophysics, Faculty of Biochemistry, Biophysics and Biotechnology, Jagiellonian University, Kraków, Poland; Sano Centre for Computational Medicine, Kraków, Poland; Department of Computers, Informatics and Microelectronics, Technical University of Moldova, Chisinau, 2045, Moldova; Department of Biological and Morphofunctional Sciences, College of Medicine and Biological Sciences, Stefan cel Mare University of Suceava, Suceava, 720229, Romania; Małopolska Centre of Biotechnology, Jagiellonian University, Kraków, Poland; EduPath, Los Angeles, CA, USA; Department of Electrical Engineering and Computer Science, Stefan cel Mare University of Suceava, Suceava, 720229, Romania; Department of Computer Science, College of Arts and Sciences, Georgia State University, Atlanta, GA, USA; Department of Systems and Computational Biomedicine, Weill Cornell Medicine, New York, NY, 10065, USA; WorldQuant Initiative for Quantitative Prediction, Weill Cornell Medicine, New York, NY, 10065, USA; Computer Science Department, Kyiv School of Economics, Kyiv, Ukraine; Medical Technologies Center, LLC “CIVITTA UKRAINE”, Kyiv, Ukraine; NGO “Genetically Modified Organisation”, Kyiv, Ukraine; Department of Biology, Uzhhorod National University, Uzhhorod, 88000, Ukraine; Broad Clinical Labs, Cambridge, MA, 02142, USA; Broad Institute of MIT and Harvard, Cambridge, MA, 02142, USA; Thomas F. Frist, Jr. College of Medicine, Belmont University, TN, 37212, USA; Genome Competence Center (MF1), Method Development and Research Infrastructure, Robert Koch Institute, Berlin, 13353, Germany; Department of Data Science and Engineering, Silesian University of Technology, Gliwice, Poland; Department of Software Engineering and Automation and Department of Industrial and Product Design, Technical University of Moldova, Chisinau, 2045, Moldova; Department of Computers, Electronics and Automation, Stefan cel Mare University of Suceava, Suceava, 720229, Romania; Sage Bionetworks, Seattle, WA, USA; Department of Computers, Informatics, and Microelectronics, Technical University of Moldova, Chisinau, 2045, Moldova; Department of Clinical Pharmacy, Alfred E. Mann School of Pharmacy and Pharmaceutical Sciences, University of Southern California, Los Angeles, CA 90089, USA

## Abstract

Disparities in formal bioinformatics training exacerbate the global skills gap, impeding the democratized application of advanced genomic technologies. To bridge this divide, we introduce a scalable, hybrid training framework designed to rapidly accelerate regional bioinformatics capacity. We exemplify this approach through the Eastern European Bioinformatics and Genomics (EEBG) workshop series — a cross-disciplinary initiative that pairs international faculty with local institutions to deliver modular, hands-on curricula. Functioning as a structured knowledge-transfer pipeline, the series has catalyzed a sustainable educational ecosystem, evidenced by the establishment of multiple independent summer schools across the region. The assessment of the 2025 EEBG workshop in Kraków, Poland, validates the model’s viability; participant metrics confirm high efficacy in skill acquisition (mean satisfaction: 4.4/5.0) and community building. Crucially, the hybrid delivery mode dismantled geographic barriers, serving as a vital mechanism for maintaining scientific continuity for researchers facing displacement and crisis. Synthesizing these outcomes, we define the core features of a replicable blueprint for scientific readiness in resource-constrained environments. We conclude by presenting a strategic roadmap — organized around infrastructure standardization, governance sustainability, and geographical expansion — for adapting this regional proof-of-concept into a global export-ready model, offering a critical path toward ensuring universal access to genomic innovation.

## Introduction

Modern life-science research has shifted toward a data-intensive paradigm, making bioinformatics skills essential across biology and medicine^1^. Yet, a significant gap persists globally between the generation of high-throughput data and the ability to analyze it. Core computational and analytical competencies remain underrepresented in life-science education programs, leaving a considerable gap between theoretical biological knowledge and practical bioinformatics proficiency^2^. Consequently, many scientists must acquire these skills outside of traditional academic settings. Global analyses reveal an overwhelming demand for short courses to build competency in biomedical data analysis and interpretation, with calls to offer such training earlier in scientists’ careers^3^.

In the Central and Eastern Europe (CEE) region, the training deficit is compounded by significant infrastructural and economic stratification. The region is not monolithic; it encompasses high-income economies like Poland and the Slovak Republic alongside lower-middle-income nations such as Moldova and Ukraine. However, cross-national indicators reveal persistent disparities in disposable income and resource distribution, notably in Romania and Bulgaria, where internal regional inequalities remain pronounced. Consequently, while some institutions enjoy modern facilities, many researchers across the region still face hurdles in accessing consistent, specialized mentorship. This uneven landscape creates a fragmented scientific ecosystem where the demand for advanced ‘omics’ analysis frequently exceeds the local supply of qualified instructors.

To mitigate these global skill deficits, international consortia such as GOBLET^4^, H3ABioNet^5^, and The Carpentries^6^ have mobilized to standardize curricula and cultivate instructor communities. Yet, the reach of these global networks has been uneven. Central and Eastern Europe has historically remained on the periphery of these initiatives, necessitating more targeted, region-specific interventions. Consequently, the establishment of formal bioinformatics education — such as dedicated degree programs and doctoral tracks— is a comparatively recent development in CEE. This delay has created a ‘critical mass’ problem: without a foundational generation of local experts, institutions struggle to staff the very programs needed to close the gap^7^.

Addressing these educational bottlenecks requires solutions that are both scalable and sustainable^8^. A successful framework combines two complementary strategies: distributed, open collaboration models that ensure continuity despite infrastructural limits, and networked training that democratizes access to cutting-edge methodologies. Within this ecosystem, targeted and intensive hybrid workshops serve as a critical bridge. By decoupling participation from geography, the hybrid format eliminates traditional administrative and travel barriers, significantly broadening accessibility for researchers in underserved regions^9^. Beyond delivering conceptual foundations and hands-on experience with real datasets, these workshops provide participants with direct networking opportunities with international mentors and peers. This approach seeds a regional community of practice that persists beyond the event. Such initiatives have proven highly effective in fostering bioinformatics capacity in resource-constrained regions, as evidenced by the establishment of annual programs across the African continent^10^.

In this work, we present a comprehensive framework for strengthening bioinformatics capacity, leveraging the Eastern European Bioinformatics and Genomics (EEBG) workshop series as a paradigmatic case study. By synthesizing lessons from this regional proof of concept, we outline a strategic roadmap for adapting this hybrid model to other underserved ecosystems. We conclude by outlining essential policy and funding mechanisms required to ensure long-term sustainability, defining a critical path toward democratizing access to genomic expertise worldwide.

### Establishing Bioinformatics Programs Helps Diversify Scientific Talent

Strategic investment in bioinformatics serves as a powerful lever for diversifying and strengthening the global scientific community. To be effective, capacity-building must simultaneously cultivate advanced expertise and democratize access to these critical skills. Prioritizing underserved academic ecosystems — particularly in Central and Eastern Europe (CEE) — accelerates the development of high-skilled STEM professionals while fostering a more inclusive research landscape. These programs yield immediate dividends: they establish a local workforce equipped with globally relevant competencies, creating modern career pathways in regions where STEM opportunities are historically limited. By mitigating ‘brain drain’ and encouraging talent retention, such initiatives generate significant socio-economic value^11^, a model successfully demonstrated by the Bioinformatics Community PL (BCPL) initiative^12^. Ultimately, local investment in bioinformatics not only reduces unemployment and elevates institutional prestige but also ensures that the global knowledge economy is driven by resilience and opportunity.

### Strategic Models for Sustainable Bioinformatics Capacity Building

To effectively address the demand for advanced bioinformatics training in resource-constrained regions, sustainable models must couple intensive, interdisciplinary instruction with direct access to computational infrastructure and cloud-based resources. This dual approach enables participants to bridge the gap between theoretical fundamentals and the practical application of high-throughput data analysis. Beyond technical skill acquisition, this format facilitates direct engagement with global experts, catalyzing the formation of a lasting regional bioinformatics ecosystem. The Eastern European Bioinformatics and Computational Genomics (EEBG) workshop series (https://eebg2025.edu.pl/) exemplifies this strategy. Designed to enable scientists in developing research environments to conduct high-impact research, EEBG combines modular training with mentorship networks. The initiative was originally conceived and launched at the Technical University of Moldova in Chişinău, which hosted the first two editions of the workshop in 2022 and 2023, laying the foundations for the EEBG network and its regional research community. Operating on a rotating host paradigm to maximize its regional footprint, EEBG subsequently expanded to the University of Suceava in Romania in 2024, before reaching the Jagiellonian University in Kraków, Poland. The 2025 iteration in Kraków, supported by the Małopolska Centre of Biotechnology and Oakland University (USA), leveraged its institutional stability to reinforce professional networking. Crucially, to ensure broad inclusion, EEBG partners with local bodies such as the Jagiellonian University Student Association ‘In Silico’ and collaborates with Bioinformatics for Ukraine to facilitate the participation of Ukrainian researchers^13^.

The EEBG consortium, comprising faculty and organizers from all past editions, advances three strategic objectives designed for global transferability. First, capacity building empowers local laboratories to independently utilize open, cutting-edge workflows. Second, talent retention connects students with international mentor networks, facilitating professional growth and global scientific integration without necessitating physical migration. Third, research continuity fosters cross-border collaborations that endure well beyond the workshop’s conclusion. Central to achieving these goals is a ‘rolling mentorship’ model, which transforms graduates into future instructors and regional capacity builders. Equipped with transferable curricula and cloud-ready workflows, EEBG alumni actively seed new networks — organizing study groups, maintaining shared digital communities, and launching satellite workshops. This multiplier effect is evident in the establishment of multiple alumni-led summer schools, particularly in Ukraine, including the Ukrainian Biological Data Science Summer School (UBDS3)^14^ in Uzhhorod and the Summer School on Computational Research Advances in Biomedical Sciences (CRABS) in Kyiv^15^. Additionally, the successful launch of the 2024 EEBG Satellite Summer School in Moldova demonstrates the model’s replicability. These regional hubs serve as powerful proof that participants can effectively transfer knowledge and adapt reusable tools to new settings, significantly expanding the model’s reach to address unmet training needs across borders^16^.

### Expanding Reach and Engagement through Hybrid Training

The fourth annual EEBG Workshop, held from July 7 to 12, 2025, at the Małopolska Centre of Biotechnology (MCB) of Jagiellonian University in Kraków, marked a significant evolution in the program’s delivery. Leveraging the advanced infrastructure of the MCB (https://mcb.uj.edu.pl/en_GB), this edition implemented a fully hybrid training architecture, offering synchronized on-site and online participation for the first time. The selection process prioritized candidates with strong biological or computational foundations who demonstrated a clear need to apply advanced genomics within their immediate professional practice. To ensure equitable learning outcomes across both cohorts, we deployed a stratified technical support architecture designed to mitigate the communication bottlenecks inherent to hybrid instruction (Box 1).

Data from the 2025 edition reveals that the hybrid format effectively bifurcated the workshop’s impact, allowing it to serve two distinct strategic functions simultaneously. Consistent with the consortium’s regional mission, the on-site cohort served as a hyper-local training hub, with 98% of in-person attendees originating from Poland (72%) and Ukraine (26%), thereby facilitating deep community building among neighbors. In contrast, the remote option acted as a global democratization engine. While overall regional participation remained robust — with 82% of all participants hailing from Poland and Ukraine — the online modality successfully dissolved geographic barriers for the remaining 18%. This remote track attracted scholars from 18 distinct nations across four continents (including Brazil, India, Kenya, and the USA), proving that hybrid architectures can maintain a strong regional core while simultaneously offering a scalable ‘on-ramp’ for the scalable ‘on-ramp’ for the Global South (Figure 1a-c; Tables S1-S2)

**Figure 1.**
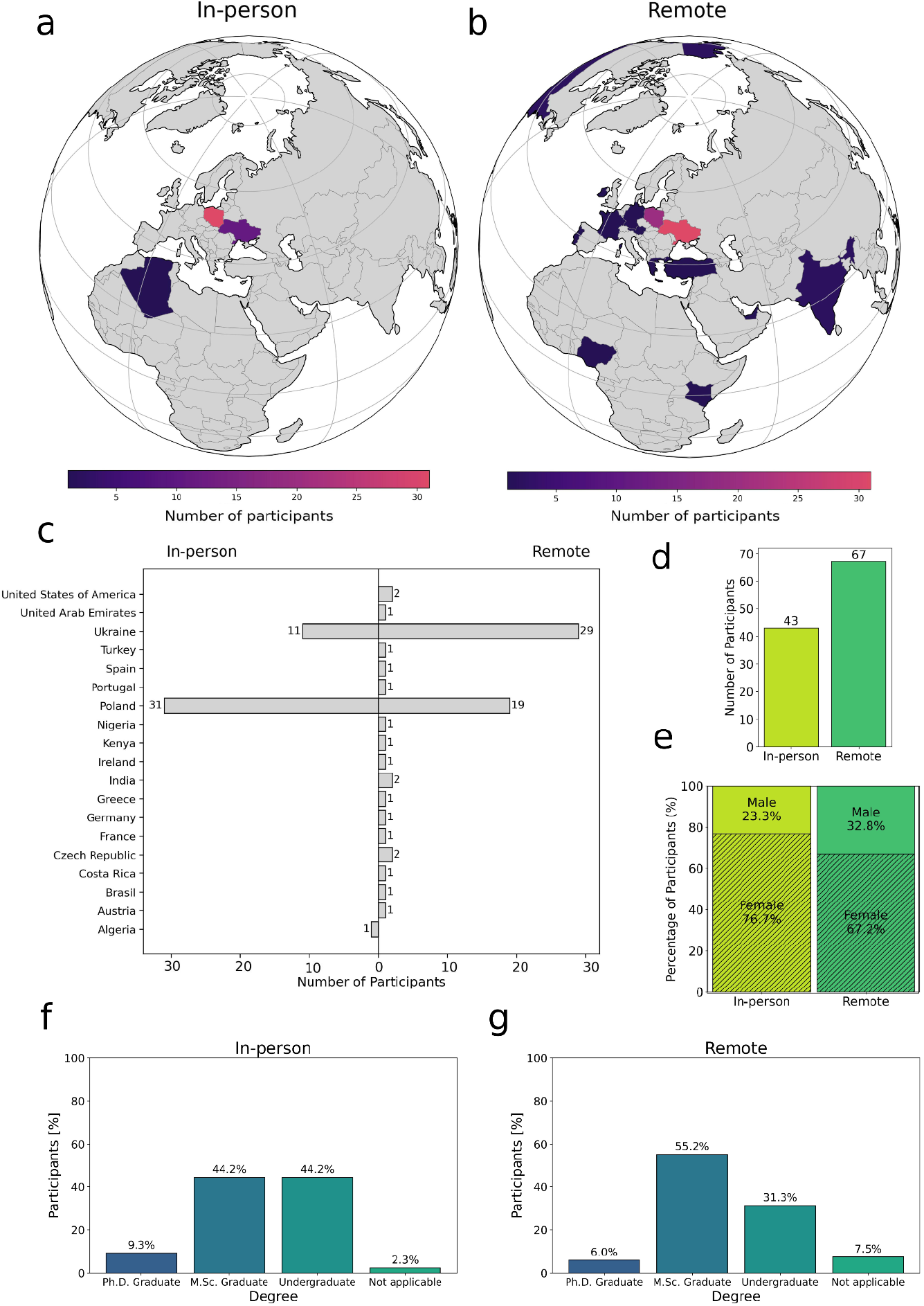
Geographic and demographic distribution of EEBG Workshop 2025 participants. (a and b) Geographic distribution of the countries of residence for participants attending (a) in person and (b) remotely. The color scale indicates the number of participants per country (purple to yellow: 0 to >30).(c) Number of in-person (left) and remote (right) participants stratified by country of residence. (d and e) Participation statistics by mode. (d) Total number of participants by attendance mode. (e) Gender distribution by attendance mode (in-person: yellow-green, left; remote: dark green, right). Diagonal hatching denotes female participants; solid fill denotes male participants. (f and g) Educational background. Survey respondents (n=45) were characterized by field of study (f) and highest academic degree obtained (graduate level) (g).

A key insight from the demographic analysis is the shifting balance between participation and physical mobility (Figure 1d). Although female participants constituted the majority in both cohorts, the proportion of male attendees was noticeably higher in the remote group compared to the on-site group (Figure 1e). This shift is particularly relevant when analyzing the Ukrainian sub-cohort (Figure S2), where martial law restricts the movement of men aged 18-60. The data suggest that while the workshop remained female-led, the hybrid format provided a critical access mechanism for male researchers who were otherwise structurally excluded from on-site attendance.

Beyond regional accessibility, a core objective of the 2025 edition was to foster cross-disciplinary engagement by bridging the divide between experimental and computational science. Survey data confirms the successful realization of this goal: the cohort was effectively split between ‘wet-lab’ biologists explicitly seeking computational upskilling (50%) and participants with established backgrounds in bioinformatics (34%), computer science/AI (14%), and exact sciences (2%). This diversity created a unique peer-learning environment. Admission protocols prioritizing candidates with pre-existing coding familiarity resulted in a distinct distribution: computationally oriented participants were significantly more likely to attend on-site (81%), whereas wet-lab biologists were evenly split between modalities (Figure S1a).

In terms of career stage, the cohort consisted predominantly of MSc graduates, complemented by smaller groups of undergraduates and doctoral candidates (Figure 1f–g). This demographic profile confirms that the curriculum successfully reached its primary target: early-career biologists facing the ‘computational gap.’ By integrating these learners with peers from strong computational backgrounds, the workshop not only transferred skills but also seeded a collaborative culture essential for modern multi-disciplinary research.

### Pedagogical Framework and Validated Learning Outcomes

The EEBG curriculum is anchored in a “didactic-to-applied” pedagogical model designed to bridge the gap between conceptual understanding and technical execution. The daily schedule utilized a dual-phase structure: morning plenary sessions established the theoretical and computational foundations, which were immediately reinforced during afternoon hands-on laboratories. This format ensured that participants not only grasped the underlying logic of genomic algorithms but also acquired the practical proficiency to implement them. The 2025 syllabus was structured around four core pillars of modern bioinformatics: genomics and metagenomics, transcriptomics, personalized medicine, and AI-assisted data science. This modular approach provided a comprehensive survey of multi-omics workflows while equipping participants with the specific tools required for independent analysis. Instruction was delivered by a cross-sector faculty, integrating academic experts from leading European and US institutions with industry specialists from Oxford Nanopore Technologies and NVIDIA, ensuring exposure to both state-of-the-art research and industrial-grade pipelines.

Post-event metrics indicate robust validation of this educational framework. Survey data (n=45; 68% in-person, 32% remote) revealed high overall satisfaction (mean 4.4 ± 0.9 /5.0; median 5.0). The didactic component received particularly high marks for clarity (mean 8.6 ± 1.4/10.00), suggesting that the theoretical scaffolding was effective. The practical sessions also scored highly (mean 8.2 ± 2.3/10.00), though qualitative analysis identified a skill-dependent divergence in user experience. While advanced users praised the immediate relevance of the workflows, novices occasionally struggled with the steep learning curve of the command-line interface. This recurrent pattern — consistent with previous editions — points to a specific opportunity for optimization: the introduction of tiered learning tracks or pre-workshop remedial modules to homogenize technical baseline skills (Figure S1b, Table S3).

### Catalyzing a Resilient and Self-Sustaining Regional Ecosystem

The impact of the EEBG series extends well beyond the physical workshop. Each edition has successfully cultivated a persistent network of contacts, supported by shared digital infrastructure (Teams, Slack, GitHub) and post-workshop study groups. This connectivity has evolved into a self-sustaining ecosystem where peer-to-peer collaborations mature into joint data analyses, cross-institutional research proposals, and international mentorship exchanges. The momentum is evident in the high rate of participant engagement: alumni frequently express a desire to return or to replicate the model by organizing satellite computational biology events at their home institutions. This dynamic signals the emergence of a distinct regional bioinformatics community — one that begins with shared technical interests but matures into a professional cohort capable of independent scientific development.

Crucially, this interconnectedness cultivates a resilient Ukrainian bioinformatics community capable of operating under conditions of displacement and crisis^17^. This objective aligns with the core mission of the Bioinformatics for Ukraine initiative, which has been integrated into the EEBG framework since 2022. By providing open-access workflows, cloud-ready computational solutions, and direct links to international mentors, the EEBG consortium facilitates the sustained integration of Ukrainian researchers into the global scientific ecosystem. This support is vital as local research infrastructure remains under severe strain due to the infrastructural instability caused by the ongoing war^13^. Ultimately, the EEBG model transcends traditional training; it is a vital instrument for maintaining scientific continuity under crisis conditions.

### Recommendations for Sustaining Global Bioinformatics Training

The organization of the EEBG series has revealed that while hybrid formats offer transformative potential for democratization, they introduce distinct logistical complexities. Delivering high-quality instruction across borders requires a rigorous operational framework that goes beyond standard workshop planning. To sustain and expand this model, organizers must prioritize technical resilience. Seamless audio-visual integration is non-negotiable; this requires investment in dedicated hybrid-classroom hardware to prevent friction. Furthermore, to maximize instructional time, we recommend a strict protocol of pre-emptive technical onboarding, where virtual machine (VM) images, datasets, and installation scripts are distributed and tested at least two weeks prior to the event. This preparation, combined with the “stratified support” system detailed in Box 1, ensures that live sessions remain focused on scientific inquiry rather than software troubleshooting.

Pedagogically, the intensity of hybrid learning demands structural adaptation to manage participant cognitive load. Future iterations should address the skill-dependent divergence observed in participant feedback by implementing tiered practical sessions — offering distinct ‘foundational’ and ‘advanced’ tracks— or mandating a pre-workshop “Zero-Day” boot camp to homogenize baseline proficiency in command-line environments. Additionally, the daily schedule must be optimized for remote engagement; instituting shorter, more frequent breaks is essential to mitigate screen fatigue and maintain high cognitive performance throughout the day.

Finally, the focus must shift from ephemeral training to an enduring community. The informal network of Slack and GitHub channels established during the workshop should be professionalized into a structured mentorship network. We recommend that future organizers formalize these interactions into an alumni ecosystem, ideally supported by a micro-granting mechanism. This financial scaffolding is crucial for enabling participants to transition from passive learners to active collaborators, funding follow-up projects and satellite events that extend the workshop’s impact into their home institutions.

### A Strategic Roadmap for Global Scaling

Building on the proven efficacy of the EEBG pilot, our future work focuses on transmuting this regional success into a standardized, globally scalable framework. We have developed a comprehensive roadmap to address the critical systemic gaps identified in recent literature^8^, specifically targeting curriculum consistency, mentorship sustainability, and resource democratization. This roadmap, detailed in Box 2, is organized into three strategic pillars — infrastructure, sustainability, and expansion — designed to embed the EEBG model into the global scientific ecosystem.

To establish the infrastructure and standardization pillar, the immediate priority is to convert our *ad hoc* successes into an export-ready asset base. To enable seamless replication, we must codify the “didactic-to-applied” method into an open pedagogical toolkit, documenting the specific technology stack and workflows required for hybrid delivery. Simultaneously, consortium leaders will work toward curriculum institutionalization by integrating EEBG content into partner university programs and establishing European Credit Transfer and Accumulation System (ECTS) accreditation pathways to validate participant effort.

The second pillar, sustainability and governance, focuses on transitioning the initiative from sporadic grant reliance to a more permanent structure. This requires securing multi-year funding to establish permanent regional hubs in the Eastern European pilot area — a strategy exemplified by the GENOMOLD program, which successfully entrenched research capacity through German-Moldovan cooperation. Crucially, long-term viability also depends on a sustainable faculty exchange program. Universities must formally recognize international mentorship as a metric for academic promotion, ensuring a continuous flow of expertise beyond initial funding cycles.

Finally, the expansion and advocacy pillar targets geographical scaling to new underserved regions, such as Southeast Asia or Latin America. This relies on a “train-the-trainer” approach, where the standardized toolkit and monitoring and evaluation (M&E) framework are deployed by new regional hosts. This expansion must be supported by vigorous policy advocacy, leveraging ROI data from the pilot phase to lobby national agencies for prioritized investment in open data infrastructures.

The feasibility of this entire ecosystem is substantially bolstered by the democratization of high-performance computing (HPC). The EuroHPC JU’s recent announcement of AI Factory Antennas (October 2025) creates a direct operational link between emerging hubs (e.g., Moldova’s FAIMA) and established compute centers (e.g., Poland’s PIAST-AI). Furthermore, the Małopolska Center of Biotechnology’s role in the Gaia AI Factory consortium — integrating with PIAST-AI and Finland’s LUMI — demonstrates how “antenna” mechanisms can distribute compute capacity. This ecosystem dramatically simplifies logistics, allowing the

EEBG model to support hundreds of concurrent users via cloud-native access programs, effectively removing hardware limitations as a barrier to global scaling.

## Discussion

The successful delivery of the Eastern European Bioinformatics and Computational Genomics 2025 workshop, synthesized with conclusions drawn from operational analysis and participant feedback, provides critical insights into the resilience of global capacity-building initiatives. The EEBG consortium’s “rolling” hybrid model validates and expands upon structures established by other regional frameworks^10^. Our results confirm that sustained, in-region training connecting local researchers with international faculty is essential for accelerating capacity development and fostering durable peer-to-peer networks. The robust participation of wet-lab biologists (approximately 50%) underscores the continuing need to embed computational training within traditional life-science disciplines — a growing necessity noted in recent literature^2,3.^ Furthermore, the hybrid delivery model effectively dismantled logistical and geographic barriers common in global bioinformatics education^8^. By lowering physical and administrative hurdles^9^, the EEBG framework demonstrates that inclusive knowledge transfer can continue even amid crisis and displacement^13^.

Beyond these structural insights, the workshop’s outcomes provide a transparent and replicable blueprint for future implementers. The “didactic-to-applied” pedagogical structure proved highly effective, achieving an average satisfaction score of 4.4/5.0. This approach enabled the rapid translation of theory into methodological independence — a critical competency for researchers in under-resourced institutions. Moreover, the spontaneous formation of alumni collaborations and study groups demonstrates that the workshop has evolved into a self-sustaining learning community, transforming a short-term training event into a long-term regional infrastructure for research continuity.

Collectively, this evidence offers compelling argumentation for engaging policymakers and funding bodies. EEBG’s success in fostering regional collaboration and ensuring scientific resilience under unstable conditions demonstrates a strong return on investment for public science funding. These results can guide policy changes that support flexible, regionally anchored training hubs and standardized curricula. Altogether, these findings provide essential recommendations for sustaining hybrid training worldwide and lay the foundation for a globally scalable roadmap organized around infrastructure standardization, governance sustainability, and strategic expansion (Box 2).

## Boxes

### Box 1. Stratified Technical Support Framework for Hybrid Delivery

Technical friction — specifically connectivity failures and environment configuration errors — constitutes the primary risk factor for participant attrition in hybrid formats. To mitigate disruption and maintain pedagogical continuity across physical and virtual cohorts, organizers should implement a multi-tiered support architecture:

- **Centralized Administrative Channel (Public):** Establish a single, high-visibility digital channel (e.g., Slack #general) dedicated exclusively to logistical directives — schedules, Zoom links, connectivity credentials, and global announcements. This “broadcast” approach ensures that administrative resolutions are immediately visible to the entire cohort, reducing redundant queries.
- **Module-Specific Technical Channels (Public):** Create distinct channels corresponding to specific curriculum modules (e.g., #genomics, #metagenomics). This segregates technical troubleshooting by topic, ensuring that discussions remain contextually relevant and allowing participants to locate solutions for specific algorithmic or software issues rapidly.
- **Tiered Escalation to 1-on-1 Support (Private):** Implement a protocol for escalating persistent or complex configuration failures to direct messaging. While public troubleshooting benefits the group, issues involving sensitive data (e.g., account credentials) or those requiring extensive debugging should be moved to private channels to prevent “noise” in the public forum.
- **Specialized Teaching Assistant (TA) Allocation:** Assign TAs and instructors to monitor channels that strictly match their domain expertise. This specialization reduces response latency and ensures that technical guidance is methodologically accurate.
- **Pre-emptive Technical Onboarding:** To maximize instructional time, shifting the “setup phase” to a pre-workshop window is critical. Virtual machine (VM) images, installation scripts, and datasets must be distributed at least two weeks prior to the event, allowing the support team to resolve compatibility issues before the live sessions commence.

### Box 2. Strategic Roadmap for Global Bioinformatics Capacity

#### Pillar 1: Infrastructure & Standardization

- ***Open Pedagogical Toolkit:*** *Develop a modular, open-source repository of workflows, teaching materials, and cloud infrastructure guides to ensure low-friction replication. (Stakeholders: Educational Technologists)*
- ***Curriculum Institutionalization:*** *Embed EEBG workshops into partner university programs (e*.*g*., *European Credit Transfer and Accumulation System credits) to ensure academic recognition and continuity between workshops and formal education. (Stakeholders: Academic Institutions)*
- ***Impact Monitoring:*** *Implement a unified M&E framework to track participant learning outcomes, career progression, and collaborative outputs over long horizons. (Stakeholders: Consortium Implementers)*

#### Pillar 2: Sustainability & Governance

- ***Regional Hub Development:*** *Secure multi-year funding to establish permanent, rotating bioinformatics training nodes across the target region. (Stakeholders: Policymakers, Funding Agencies)*
- ***Local Ownership Transfer:*** *Transition governance of regional hubs to local institutions to ensure long-term autonomy while maintaining global alignment. (Stakeholders: Local Host Institutions)*
- ***Sustainable Faculty Exchange:*** *Formalize agreements where faculty service in regional workshops is recognized in academic promotion and tenure tracks. (Stakeholders: University Leadership)*

#### Pillar 3: Expansion & Advocacy

- ***Pilot Expansion:*** *Launch the standardized model in new underserved regions (e*.*g*., *Southeast Asia, Latin America) using the established toolkit and mentorship framework. (Stakeholders: Global Funding Bodies)*
- ***Seed Funding Mechanism:*** *Create a competitive micro-grant program for alumni-led research projects and satellite workshops to foster scientific independence. (Stakeholders: Funding Agencies, Alumni Committee)*
- ***Policy Advocacy:*** *Leverage ROI data from the pilot phase to lobby national and international agencies for prioritized investment in open data infrastructures. (Stakeholders: Policymakers, nonprofit organizations)*

