## Supplementary Tables and Figures for "Leveraging a hybrid cross-disciplinary training model to accelerate global bioinformatics capacity"

**Table S1. Country of residence of EEBG 2025 applicants by attendance mode.** Number (n) and percentage (%) of applicants attending in person or remotely are shown for each country, together with overall totals.

| Country of Residence | In-person (n) | In-person (%) | Remote (n) | Remote (%) | Overall (n) | Overall (%) |
|---|---|---|---|---|---|---|
| Algeria | 1 | 1,28 % | 0 | - | 1 | 0,78 % |
| Australia | 1 | 1,28 % | 0 | - | 1 | 0,78 % |
| Austria | 0 | - | 1 | 1,96 % | 1 | 0,78 % |
| Brasil | 0 | - | 1 | 1,96 % | 1 | 0,78 % |
| Colombia | 1 | 1,28 % | 0 | - | 1 | 0,78 % |
| Costa Rica | 0 | - | 1 | 1,96 % | 1 | 0,78 % |
| Czech Republic | 1 | 1,28 % | 1 | 1,96 % | 2 | 1,55 % |
| Estonia | 1 | 1,28 % | 0 | - | 1 | 0,78 % |
| France | 0 | - | 1 | 1,96 % | 1 | 0,78 % |
| Germany | 3 | 3,85 % | 0 | - | 3 | 2,33 % |
| Greece | 0 | - | 1 | 1,96 % | 1 | 0,78 % |
| India | 1 | 1,28 % | 1 | 1,96 % | 2 | 1,55 % |
| Ireland | 0 | - | 1 | 1,96 % | 1 | 0,78 % |
| Kenya | 0 | - | 1 | 1,96 % | 1 | 0,78 % |
| Nigeria | 1 | 1,28 % | 1 | 1,96 % | 2 | 1,55 % |
| Pakistan | 1 | 1,28 % | 0 | - | 1 | 0,78 % |
| Poland | 42 | 53,85 % | 16 | 31,37 % | 58 | 44,96 % |
| Portugal | 0 | - | 1 | 1,96 % | 1 | 0,78 % |
| Spain | 0 | - | 1 | 1,96 % | 1 | 0,78 % |
| Turkey | 0 | - | 1 | 1,96 % | 1 | 0,78 % |
| Ukraine | 25 | 32,05 % | 19 | 37,25 % | 44 | 34,11 % |
| United Arab Emirates | 0 | - | 1 | 1,96 % | 1 | 0,78 % |
| United States of America | 0 | - | 2 | 3,92 % | 2 | 1,55 % |
| **Total** | **78** | **100,00 %** | **51** | **100,00 %** | **129** | **100,00 %** |

**Table S2. Country of residence of EEBG 2025 participants by attendance mode.** Number (n) and percentage (%) of participants attending in person or remotely are shown for each country, together with overall totals.

| Country of Residence | In person (n) | In person (%) | Remote (n) | Remote (%) | Overall (n) | Overall (%) |
|---|---|---|---|---|---|---|
| Algeria | 1 | 2,33 % | 0 | - | 1 | 0,91 % |
| Austria | 0 | - | 1 | 1,49 % | 1 | 0,91 % |
| Brasil | 0 | - | 1 | 1,49 % | 1 | 0,91 % |
| Costa Rica | 0 | - | 1 | 1,49 % | 1 | 0,91 % |
| Czech Republic | 0 | - | 2 | 2,99 % | 2 | 1,82 % |

| | | | | | | |
|---|---|---|---|---|---|---|
| France | 0 | - | 1 | 1,49 % | 1 | 0,91 % |
| Germany | 0 | - | 1 | 1,49 % | 1 | 0,91 % |
| Greece | 0 | - | 1 | 1,49 % | 1 | 0,91 % |
| India | 0 | - | 2 | 2,99 % | 2 | 1,82 % |
| Ireland | 0 | - | 1 | 1,49 % | 1 | 0,91 % |
| Kenya | 0 | - | 1 | 1,49 % | 1 | 0,91 % |
| Nigeria | 0 | - | 1 | 1,49 % | 1 | 0,91 % |
| Poland | 31 | `72,09 % | 19 | 28,36 % | 50 | 45,45 % |
| Portugal | 0 | - | 1 | 1,49 % | 1 | 0,91 % |
| Spain | 0 | - | 1 | 1,49 % | 1 | 0,91 % |
| Turkey | 0 | - | 1 | 1,49 % | 1 | 0,91 % |
| Ukraine | 11 | 25,58 % | 29 | 43,28 % | 40 | 36,36% |
| United Arab Emirates | 0 | - | 1 | 1,49 % | 1 | 0,91 % |
| United States of America | 0 | - | 2 | 2,99 % | 2 | 1,82 % |
| **Total** | **43** | **100 %** | **67** | **100 %** | **110** | **100 %** |

**Table S3.** The EEBG 2025 Workshop assessments by participants depending on the participation mode. The table contains mean values with standard deviations; the scale of assessment was 1 to 10.

| | **In-person** | | **Remote** | |
|---|---|---|---|---|
| | **Lectures** | **Hands-On Workshops** | **Lectures** | **Hands-On Workshops** |
| Day 1: Frontiers in Computational Biology | 8.20 ± 1.39 | 9.10 ± 1.45 | 8.82 ± 1.38 | 7.64 ± 2.68 |
| Day 2: Metagenomics | 8.29 ± 1.42 | 6.67 ± 2.83 | 9.02 ± 1.25 | 8.43 ± 2.38 |
| Day 3: Personalized Medicine | 8.76 ± 1.24 | 7.20 ± 2.54 | 8.79 ± 1.50 | 8.29 ± 2.20 |
| Day 4: Oxford Nanopore Technology | 8.17 ± 1.78 | 8.23 ± 1.85 | 8.98 ± 1.38 | 7.86 ± 1.88 |
| Day 5: Transcriptomics | 8.82 ± 1.10 | 9.60 ± 0.86 | 9.05 ± 1.07 | 9.36 ± 1.39 |
| **Overall** | **8.45 ± 1.41** | **8.16 ± 2.29** | **8.93 ± 1.28** | **8.31 ± 2.16** |

### Supplementary Figures

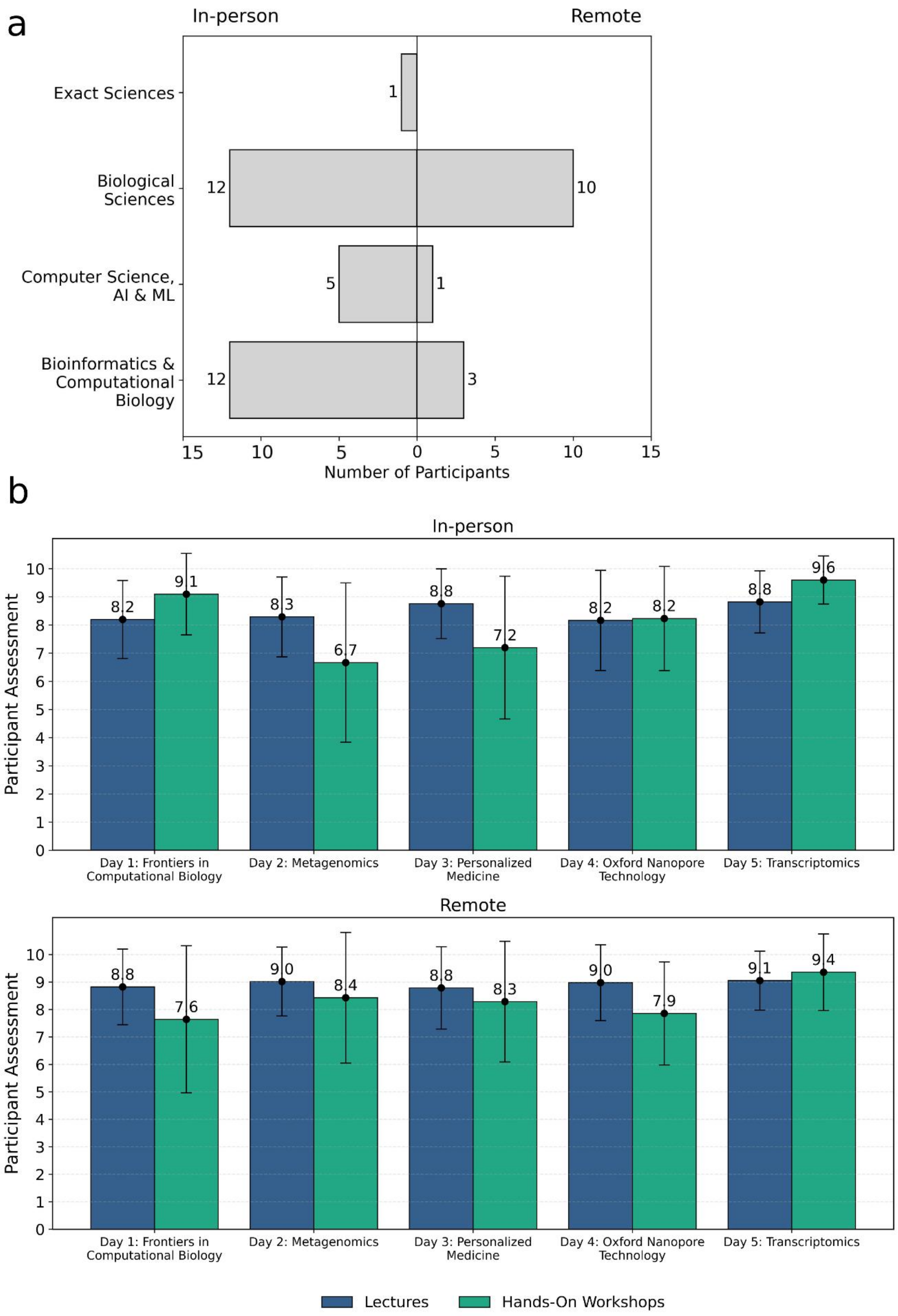

Figure S1 | (a) **Diverse educational backgrounds of EEBG 2025 workshop participants**. Number of in-person (left) and remote (right) participants by their educational/research background: exact sciences, biological sciences, computer sciences, Artificial Intelligence and Machine Learning, bioinformatics and computational biology. (b) The **overall satisfaction of the EEBG 2025 participants with the provided training depending on the participation mode.** Color represents the type of learning activity: lectures (blue) or hands-on workshops (yellow). The graph shows the overall satisfaction of the in-person (upper part, n=31) and remote (lower part, n=14) participants, who filled the survey, with the classes conducted at each of the daily thematic sections. Data are shown as mean ± standard deviation.

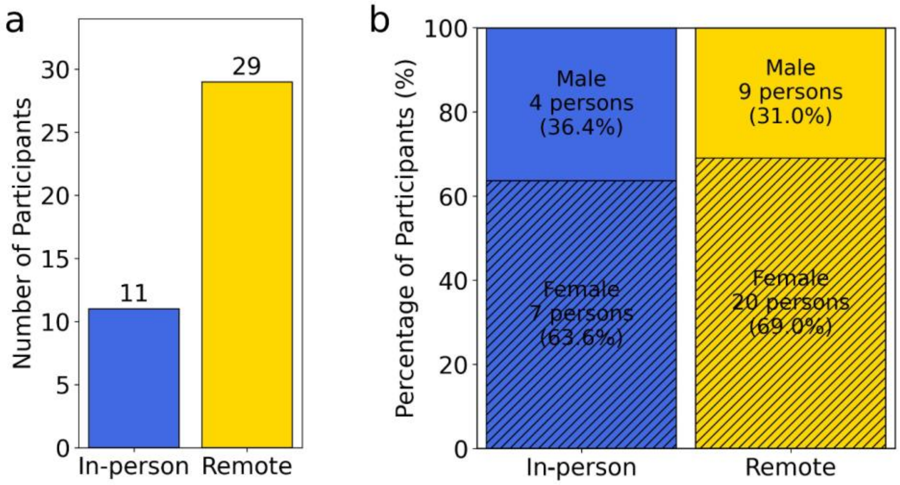


Figure S2 | **Ukrainian sub-cohort statistics by participation mode**. (a) Total number of Ukrainian participants by attendance mode, and (b) gender distribution by attendance mode (in-person: blue, left; remote: yellow, right). Diagonal hatching denotes female participants; solid fill denotes male participants.
